# The network structure of cancer ecosystems

**DOI:** 10.1101/240796

**Authors:** Simón P. Castillo, Rolando Rebolledo, Matias Arim, Michael E. Hochberg, Pablo A. Marquet

## Abstract

Ever since Paget’s seed-and-soil and Ewing’s connectivity hypotheses to explain tumor metastasis (1,2), it has become clear that cancer progression can be envisaged as an ecological phenomenon. This connection has flourished during the past two decades (*3–7*), giving rise to important insights into the ecology and evolution of cancer progression, with therapeutic implications *(8–10)*. Here, we take a metapopulation view of metastasis (i.e. the migration to and colonization of, habitat patches) and represent it as a bipartite network, distinguishing source patches, or organs that host a primary tumor, and acceptor patches, or organs colonized ultimately from the source through metastasis. Using 20,326, biomedical records obtained from literature, we show that: (i) the network structure of cancer ecosystems is non-random, exhibiting a nested subset pattern as has been found both in the distribution of species across islands and island-like habitats *(11–13)*, and in the distribution of among species interactions across different ecological networks *(14–16);* (ii) similar to ecological networks, there is a heterogeneous distribution of degree (i.e., number of connections associated with a source or acceptor organ); (iii) there is a significant correlation between metastatic incidence (or the frequency with which tumor cells from a source organ colonize an acceptor one) and arterial blood supply, suggesting that more irrigated organs have a higher probability of developing metastasis or being invaded; (iv) there is a positive correlation between metastatic incidence and acceptor organ degree (or number of different tumor-bearing source organs that generate metastasis in a given acceptor organ), and a negative one between acceptor organ degree and number of stem cell divisions, implying that there are preferred sink organs for metastasis and that this could be related to average acceptor organ cell longevity; (v) there is a negative association between organ cell turnover and source organ degree, implying that organs with rapid cell turnovers tend to generate more metastasis, a process akin to the phenomenon of propagule pressure in ecology *(17)*; and (vi) the cancer ecosystem network exhibits a modular structure in both source and acceptor patches, suggesting that some of them share more connections among themselves than with the rest of the network. We show that both niche-related processes occurring at the organ level as well as spatial connectivity and propagule pressure contribute to metastatic spread and result in a non-random cancer network, which exhibits a truncated power law degree distribution, clustering and a nested subset structure. The similarity between the cancer network and ecological networks highlights the importance of ecological approaches in increasing our understanding of patterns in cancer incidence and dynamics, which may lead to new strategies to control tumor spread within the human ecosystem.

---

In 1829, Joseph Recamier coined the term ’metastasis’ for the spread of cancer cells away from organs where a primary tumor had emerged. Despite many intervening decades of observation and investigation, our understanding of metastasis is largely centered on a small number of organs and tissues (*18,19*). These studies indicate that a series of often complex processes occur in metastasis, beginning with the migration of cancer cells from a source tumor, through numerous intermediate states, habitats and microenvironmental conditions, and culminating either in spatially distinct tumors in local tissue or in distant organs *(20,21)*. Although the basic sequence is very similar among those cancers that have been studied in detail, it is not well understood why certain organs and tissues (hereafter ’organs’) are more commonly the sites of metastasis, nor why some specific primary tumor sites tend to be associated with one or a few specific metastatic organ sites *(20)*, while others are more generalists and tend to metastasize in many different organs *(22)*.

Two main hypotheses have been offered to explain why some organs are the target for metastasis *(23)*. Under the first hypothesis (also known as the ’seed-and-soil’ hypothesis) proposed by Stephen Paget *(1)*, tumor cells migrate from established tumors and only create self-sustaining metastatic growth in distant organs if the latter’s microenvironmental conditions are adequate *(24)*. This set of conditions is analogous to the Grinellian niche in ecology *(25)*, and in cancer is the set of conditions under which cancer cells survive migration, settle and grow *(18, 26, 27)*. The second hypothesis is associated with the proposal by James Ewing’s work *(2)*, and suggests that metastatic spread occurs by purely mechanical factors associated with the anatomical structure of the vascular system. Here, the probability of an organ harboring a metastasis depends in part on the number of cancer cells delivered to it, which in turn is a function of blood flow and distance to the source organ *(28,29)*. Whereas there is little support for the sole action of the mechanical hypothesis in explaining observed patterns in metastasis, measures integrating microenvironments and mechanical variables could be more representative *(30)*.

Competing hypotheses to explain the mechanisms contributing to metastatic patterns have been evaluated with reference to the primary tumor (e.g., the lung tumor and its metastasis *(31)*), but how findings generalize across both primary and metastatic tumor sites is unknown. In this contribution, we carry out a large scale statistical analysis of metastatic pattern (using 20,326, biomedical records, see Supplementary Material) by taking a network approach of the association between an organ with a primary tumor and its metastatic sites. To do this we employ methods from metapopulation theory *(7)* and classify ’source patches’ (or *S*) as those organs where the primary tumor emerged and from where metastatic propagules migrate, and as ’acceptor patches’ (or *K*) as those organs that receive these propagules and become colonized. In this context, source organs can be associated to acceptor organs using a bipartite network or graph defined as *G* = (*S*, *K*, *E*) (Figure 1A), where source patches and acceptor patches are connected by links or ’edges’ (*E*).

**Fig. 1.**
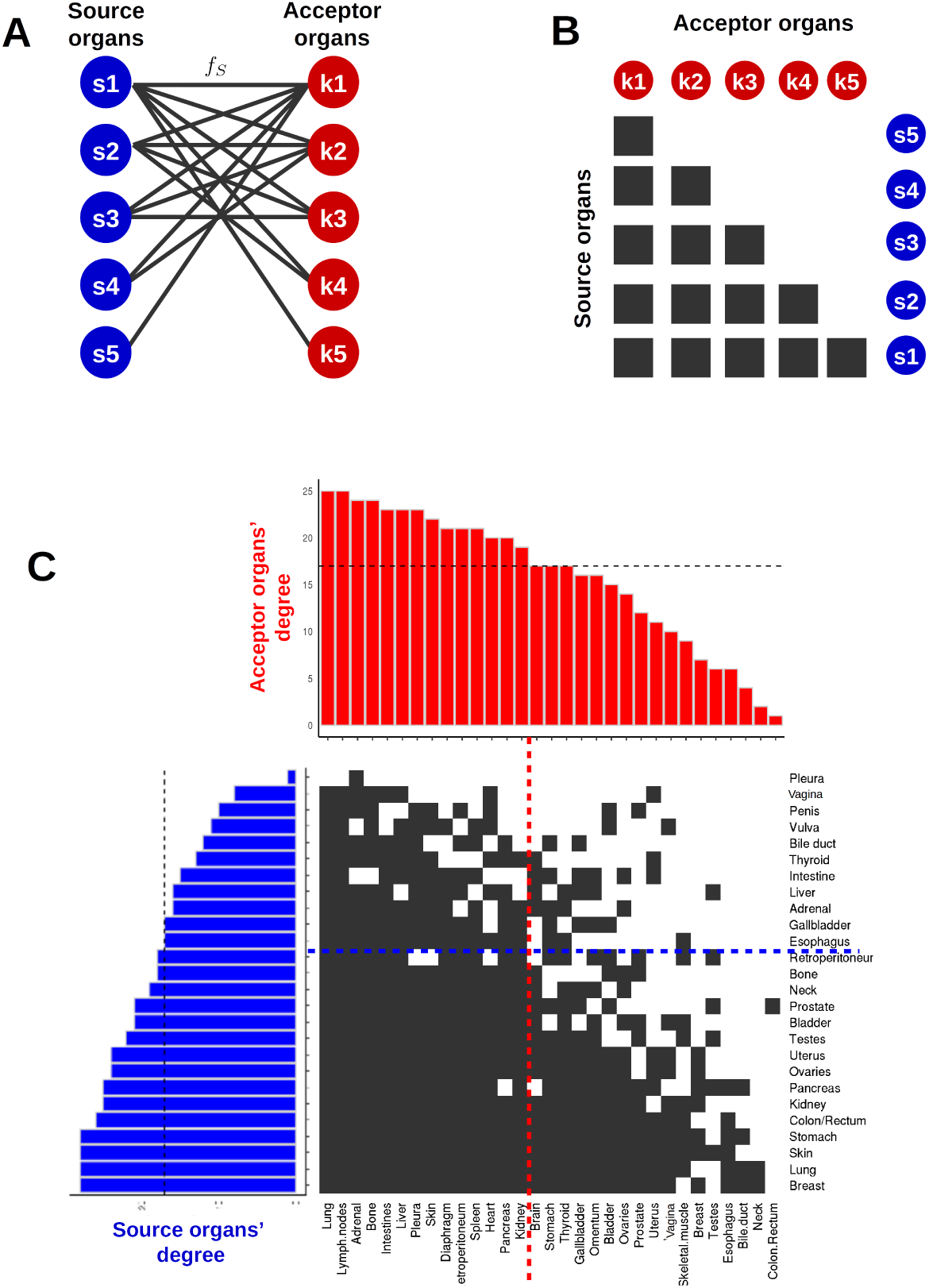
The cancer network. (A) Schematic view of the bipartite network connecting source *S* = (*S*_1_, …,*S*_5_) and acceptor *K* = *(K*_1_, …,*K*_5_) organs through metastatic propagules *f_S_: S* → *K* (B) This network can be represented as a matrix with rows and columns arranged according to their degree, in this case the matrix is perfectly nested. (C) The observed cancer matrix connecting source (rows) with acceptor (columns) organs. Histograms show the degree corresponding to each organ. Dashed lines identify the median of the degree distribution. They define a core of highly interacting organs and a periphery where low-degree organs interact with high-degree ones.

We found that source and acceptor organs vary in terms of the number of connections they have (i.e. their degree). This may be associated with a monotonic gradient in migrating cell invasiveness across source organs (i.e., some source organs are connected to more acceptor organs, and/or generate more migrating cells, and/or migrating cells that are more adapted to migration) and a monotonic gradient in invasibility across acceptor organs (i.e., some acceptor organs are more prone to be invaded than others, and/or to receive metastases from more source organs) (Figure 1C). This diversity may be associated with tissue-specific risk factors *(32, 33)*, different life history trade-offs in tumors *(34)*, and variation in the degree of matching between the quality of the recipient organ and the niche requirements of migrating metastatic cells, all of which drives colonization success (*5,24,35*), analogous to what is often observed in models of metapopulation dynamics *(7,36,37)*.

More interestingly, we found that the metastatic network is highly structured with a scalefree, truncated power-law, degree distribution that applies to both source or acceptor organs, and which is significantly different from the exponential model expected for a random network (Fig. 2, Table S1). A scale-free degree distribution is a property shared by different complex networks, from protein interactions to networks of scientific collaboration *(38–40)*. Similarly, the metastatic network is highly nested (nestedness tests: NODF = 83.12 and BINMAT = 5.95) and asymmetric, such that there is a network core of highly interacting source and acceptor organs and a periphery where specialized source (acceptor) organs interact with generalist acceptor (source) organs (Figure 1B,C). We also found that the cancer network shows clusters of interacting organs (Figure S1), which is reflected by a modular structure. This implies that there are groups of source and acceptor organs that are more similar among themselves than to the rest of the network in which they are embedded *(41,42)* as shown by *(32)* for cancer risk. Modular networks in ecology have been associated with the presence of species with similar functional traits (*43, 44*) or subsets of locations with more frequent dispersal *(45,46)*. These two mechanisms are plausible in cancer networks. Modularity may reflect a combination of similar traits among groups of organs (due either to similar organ environments or shared connectivity characteristics), and different traits between groups that restrict metastasis from certain source organ groups but not others.

**Fig. 2.**
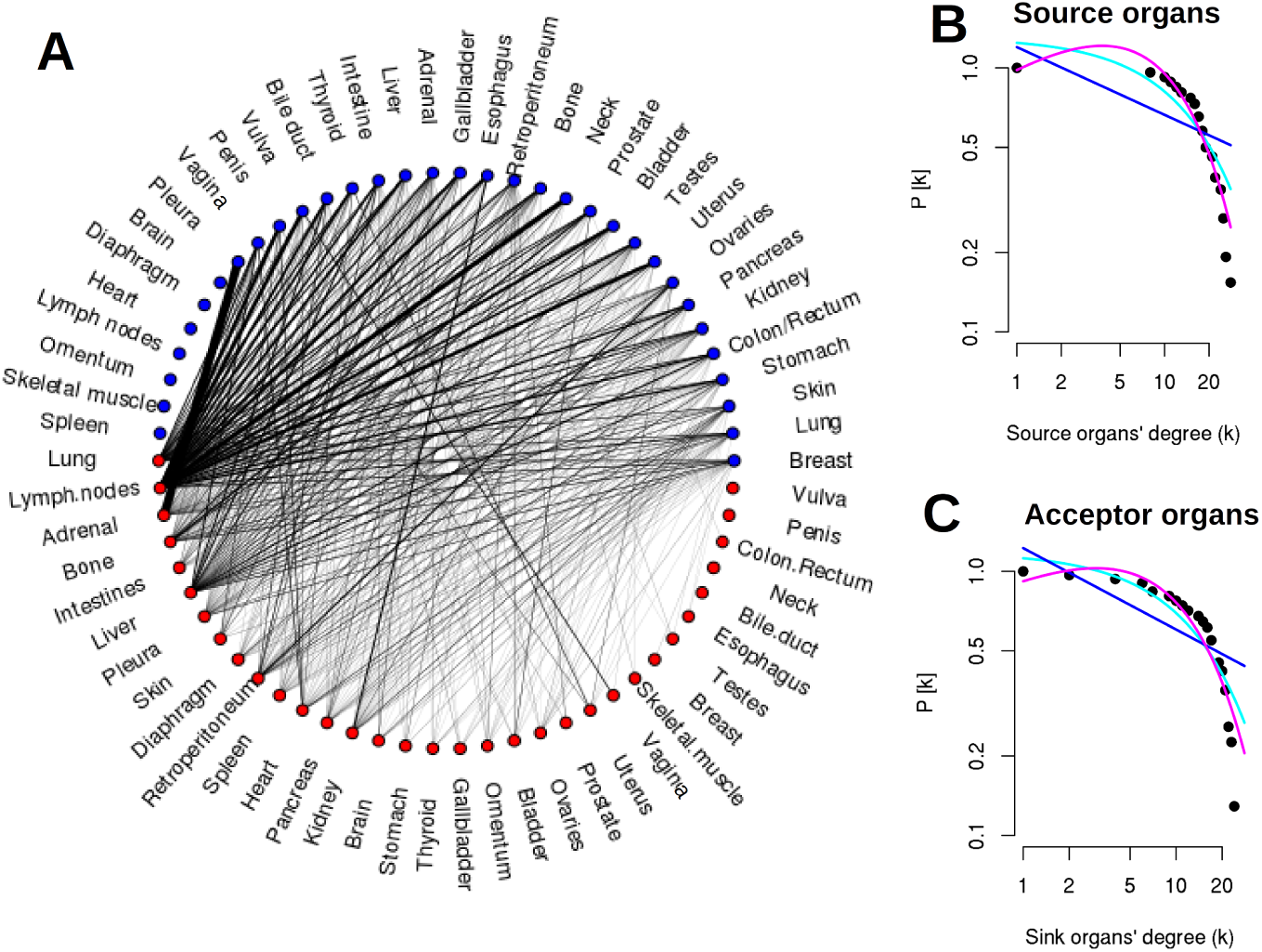
A scale-free cancer network. (A) Shows links from a source organ (blue nodes) to acceptor organs (red nodes). Links width is proportional to NSI. (B) Three different models were fitted to the probability that the degree of a randomly chosen organ is larger than k or *P*[*x* ≥ *k*] where *k* is the number of links or degree of the organ: exponential (blue), power law (cyan) and truncated-power law (magenta). The best fit model was truncated power law (for summary statistics see Table S1).

Scale-free degree distributions, modularity and nestedness patterns in networks have been suggested to promote diversity, stability and network robustness to disturbances *(14,47)*. In an oncological context, these network attributes are not related to stability and robustness, since organs are not species that can go extinct, instead our findings suggest the action of one or a few simple mechanisms *(16,39)*.

## Explaining the observed patterns

The simplest way to generate a scale free network is based on the action of two simple generic mechanisms *(39)*: 1) one that provides for the continuous increase in the number of links resulting in the expansion of the network, and 2) one that accounts for an increase in the probability of a site being connected as function of the number of connections it already has or ’preferential attachment’. For the metastatic network presented here it is important to keep in mind that it corresponds to a network reconstructed from an ensemble of cases, where each link implies that the interaction has been recorded in at least one case (i.e. an individual with a primary tumor and its corresponding metastasis). In what follows we propose that these two mechanisms can account for the network patterns in metastasis.

The first mechanism implies that the network structure has changed through time, because of carcinogenesis and cancer cell migration from novel primary sites, and/or metastasis to novel acceptor sites. Changes in tissue-level cancer risk may have an evolutionary basis (*32, 48, 49)* and/or be associated with novel environmental conditions *(50)*. The second mechanism is associated with preferential attachment, which in this context implies that a new primary tumor will likely metastasize in an acceptor organ with a high degree, and that a new metastasis is more likely to arise in a primary tumor that already metastasize to many different organs. This mechanism by definition will generate nestedness, whereby specialized (low degree) acceptor organs are more likely to interact with generalized (high degree) source organs and vice versa. The mechanism behind preferential attachment in cancer networks is likely the result of some organs being more likely to express a primary tumor as well as to receive metastases *(22)*.

As shown in Figure 3, the number of connections (i.e. its degree) that a given acceptor organ has (i.e. the number of different primary tumors that can metastasize to it), increases as the narrow sense and broad sense incidence (NSI and BSI respectively, see Supplementary Materials) of metastasis in that organ increases. Thus there are some organs that are ’preferred’ targets for metastases (hence there are many cases of these combinations in the population) from a given primary tumor (NSI) or for different primary tumors (BSI). Also, our analysis identifies a negative correlation between acceptor organ degree and the number of stem cell divisions, implying that organs which on average have fewer or older cells are targets for metastasis from a larger number of different primary tumors, in accordance with Paget’s seed and soil hypothesis. In this case, they receive more links because they are inherently more suitable to be colonized, and this is likely one of the mechanisms behind preferential attachment, and hence nestedness. Finally, the truncation phenomenon observed in the degree distribution of our networks is likely the result of the small and finite number of nodes (organs) that can potentially be part of the network *(51)*, and which limits the spread and filling of the distribution.

**Fig. 3.**
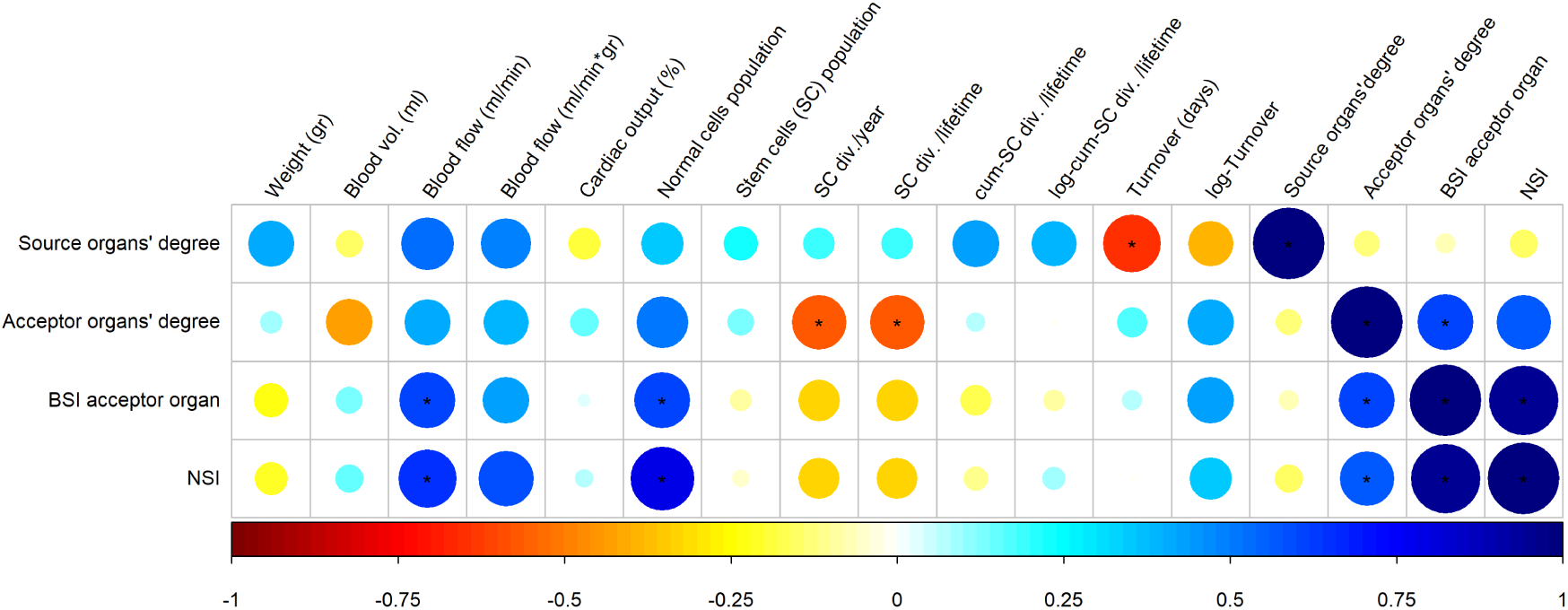
Correlation matrix testing the linear association between our network measures for each organ (rows) and data from literature (see methods for details and references). Asterisks(*) indicate that the correlation coefficient differs significantly from 0. (BSI: broad sense incidence. NSI: Narrow sense incidence).

Nestedness in ecological systems can arise because habitat patches display a gradient in either colonization or extinction probabilities (*11,52*). In the case of the cancer network presented herein, both extinction and colonization appear to influence observed patterns. Extinction in the context of metastasis corresponds to failed colonization (to the point of producing a detectable tumor), resulting from either intrinsic inhospitability of certain organs to cancer cell growth, or from characteristic non-compatibility between certain primary tumor metastatic cells and specific organ microenvironments. As per colonization, we found positive correlations between blood flow through an organ and the incidence of metastasis and this is valid for both BSI and NSI (Figure 3). Blood flow is correlated with the number of propagules that could potentially arrive in a patch, or ’propagule pressure’ *(17)*. Similarly, the degree, of a source organ is negatively correlated with cell turnover in that organ (Figure 3). This implies that source organs with more frequent cell division (i.e. shorter turnover time) generate more metastasis to different acceptor organs. Taken together, these results suggest both a role for propagule pressure, hence colonization, in the observed nested pattern of the cancer network, and the importance of stem cells’ life history and organ turnover *(53–55)* in understanding the emergence of primary tumors and metastasis. Finally, as suggested by Ewing’s hypothesis, we have found that spatial closeness related to arterial blood supply, as a proxy for spatial proximity, is correlated with metastatic incidence (Kendall-*τ* = 0.096, p-value = 0.003), suggesting that the probability of metastasis in an acceptor organ increases if it shares an artery with the source organ organ (Fig. S2) (see methods in Supporting Material).

It is clear that further research beyond a static view of network structure is needed to understand the observed variation in susceptibility of organs to metastasis. Network dynamics can be of great importance, particularly in understanding the phenomenon of tumor self-seeding *(56, 57)* or the possibility of stepping stone migration from a primary source to an acceptor organ via intermediate organs (*23, 31*). The ecological network approach presented here has the power to generate insights to direct both fundamental and applied research of therapeutic relevance. In particular, and considering the asymmetric nature of the cancer network reflected in its nested subset structure, it is important to resolve the mechanisms behind the gradient in specificity/generality observed in source and acceptor organs.

**Conclusions** We show that both niche-related processes occurring at the organ level as well as spatial connectivity and propagule pressure are consistent with patterns in metastatic spread as evidenced by a non-random cancer network, which exhibits a truncated power law behavior, clustering and nested subset structure.

## Acknowledgments

SPC acknowledges CONICYT-PFCHA/Doctorado Nacional/2017–21170089, PAM acknowl FONDECYT 116223 and CONICYT Basal Funding Program PFB-023.

## Supplementary Materials

### Materials and Methods

We studied the metastatic network between organs by constructing a bipartite network based on a matrix representing the number of occurrences that a primary neoplasm in a source organ generated a metastasis in an acceptor organ. The data was obtained from the literature *(1–12)*. Following diSibio and French *(1)*, 33 anatomical zones (referred here as ’organs’) were identified (Fig. 1). We categorized a *source* as the organ where the primary carcinoma was found, and as *acceptor* organs those organs with secondary growth of a neoplasm, according to the reported metastatic sites. A total of 20,326 occurrences were included in the analysis, based on autopsies and tomographies in the case of muscular cancers. Medical records are from the USA, Switzerland, Germany and Slovenia. In analytical terms, we define the metastatic process as a graph *G* = (*S*, *K*, *E*) where *S* and *K* denote the set of source and acceptor organs, respectively and *E* identifies the links or edges connecting them. We focus our analysis in the weighted network *(W = G = (S, K, E*)) of source-acceptor organ interactions *S_i_* × *K_j_* with *S_i_* ∈ *S*, *K_j_* ∈ *K* and *S* = *K* = (1*,…,N*), where *N* corresponds to 33 anatomical sites. Let us define *f_S_* as the metastatic process *f_S_* : *S* → *K*. The values of the source to acceptor weight (narrow sense metastatic incidence or NSI) *f_S_* ∈ [0,1] corresponds to the number of metastases found at an acceptor organ that derived from a given *source* organ out of the total number of metastases recorded for that source organ in the population of cases. Thus, *f_S_* represents the relative importance of acceptor organs for the propagules generated by the primary tumor in the source patch. Similarly, the number of times that a primary tumor was recorded in a source organ or that a metastasis was recorded in an acceptor one, out of the total number of cases in the population, corresponds to the Broad Sense Incidence or BSI of source or acceptor organs.

Vascular incidence matrices were estimated based on the main artery or vein conducting blood flow between organs. We identified the main arteries associated with an organ’s blood supply: thoracic aorta, abdominal aorta, left gastric coeliac trunk, splenic artery, common hepatic artery, superior mesenteric artery, and internal iliac artery; and those involved in blood drainage: renal veins, inferior phrenic vein, hepatic vein, gastric veins, splenic vein, mesenteric vein, internal iliac vein, and internal jugular vein. To characterize the spatial association emerging from the vascular arrangement, we constructed a vascular squared matrix *n_i_* × *n_j_* with *n* = (*n*_1_, *n*_2_, *n_i/j_,…,N*) being the number of organs (number of rows/columns). When two organs shared an artery or a vein, we recorded this co-occurrence as a 1. In contrast when a couple (*n_i_*, *n_j_*) did not share a vessel we assigned a 0. For example, because pancreas and liver share blood supply through the abdominal aorta, the supply matrix has a value of 1 associated to the pair ’pancreas, liver’. Our first approach was to correlate the weighted metastatic matrix with both vascular matrices independently, expecting that if organ connectivity plays a role in cancer spread, it will manifest in a significant statistical association. Our results show that, the drainage network is not statistically associated with metastatic incidence in source organs (Kendall-*τ* = 0.042, p-value = 0.204), although there are some cases where acceptor organs that share a proximal vein with the source have higher metastatic incidences (Fig. S3). On the other hand, the supply network, based on common arteries shared by organs, was significantly correlated with the metastatic incidence network (Kendall-*τ* = 0.096, p-value = 0.003), suggesting that organs which share an artery, on average, have higher metastatic incidences (Fig. S2). These results follows the intuitive idea that metastatic propagules leaving their source organ follow blood flow, increasing their chances of colonizing new patches.

All data analyses were performed in R *(13)*. Nestedness measures were contrasted against a random null model (function `nestedchecker`, package `vegan`) under a non-sequential algorithm for binary matrices that only preserves the number of occurrences within the matrix. In this case the statistical significance of the estimated C-score was analyzed. We tested linear correlations (null hypothesis: *ρ* = 0) of our network metrics (source organ degree, acceptor organ degree and metastatic incidence) against the following estimates extracted from the literature: From Weiss et al *(14)*: organ weight, blood volume (ml), blood flow (ml/min), mass-specific blood flow (ml/(min*gr)). From Sidhu et al 2011 *(15)*: cardiac output (%). From Tomasetti and Volgestein *(16)*: normal cell population number, number of stem cells, number of division of each stem cell per year, number of divisions of each stem cell per lifetime, cumulative divisions of each stem cell per human lifetime. From Richardson, Allan and Le *(17)*: organ turnover.

### Modularity analysis

Modularity was obtained with the package `bipartite` implementing the QuanBiMo algorithm *(18)* for bipartite networks’ module detection. This algorithm allows the detection of modules based on metastatic incidence patterns.

**Fig. S1.**
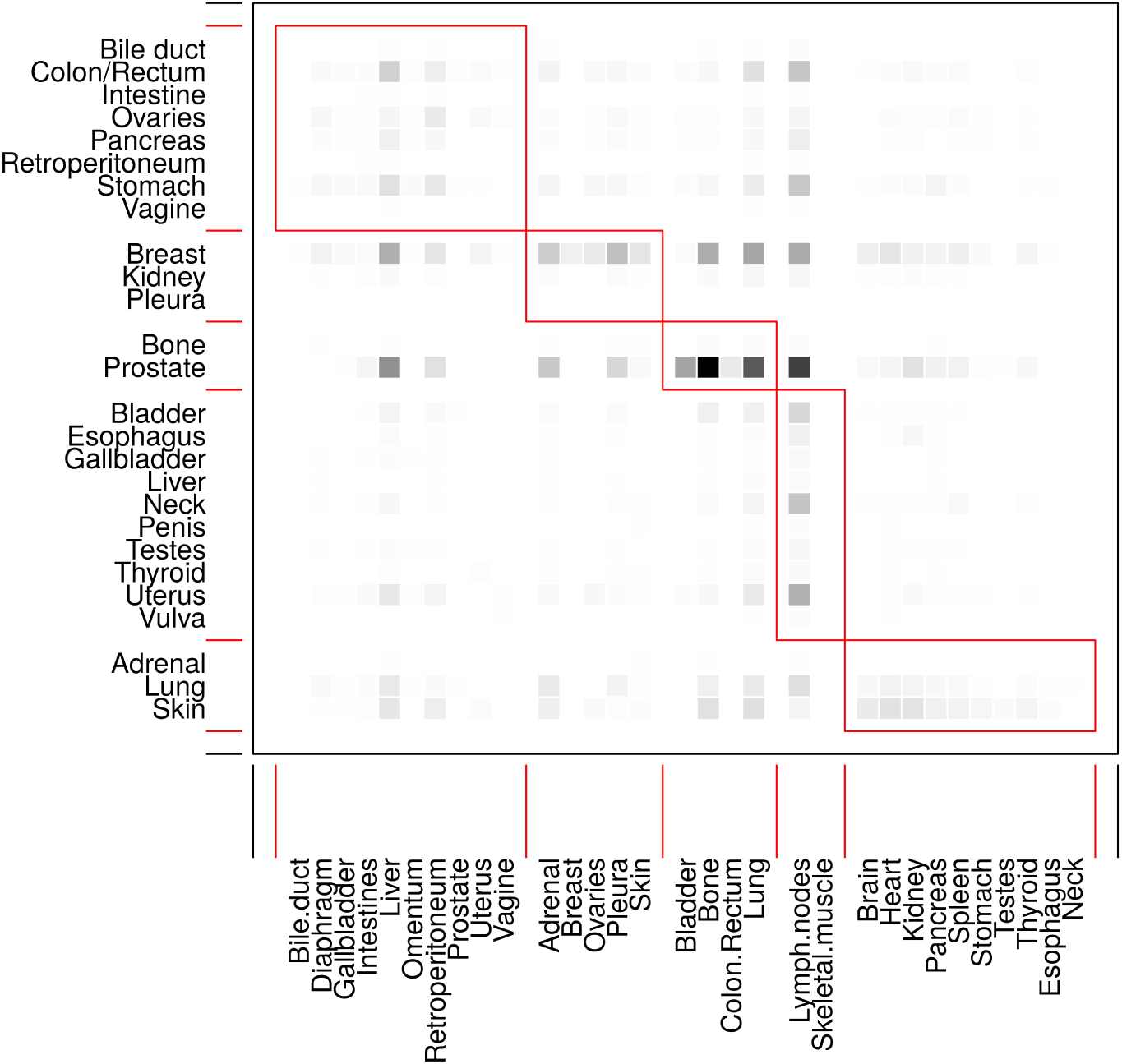
The identification of modules based on BSI for the metastatic spread from source organs (rows) to acceptor organs (columns). Red boxes delineate the modules detected by QuanBiMo algorithm *(18)*.

**Fig. S2.**
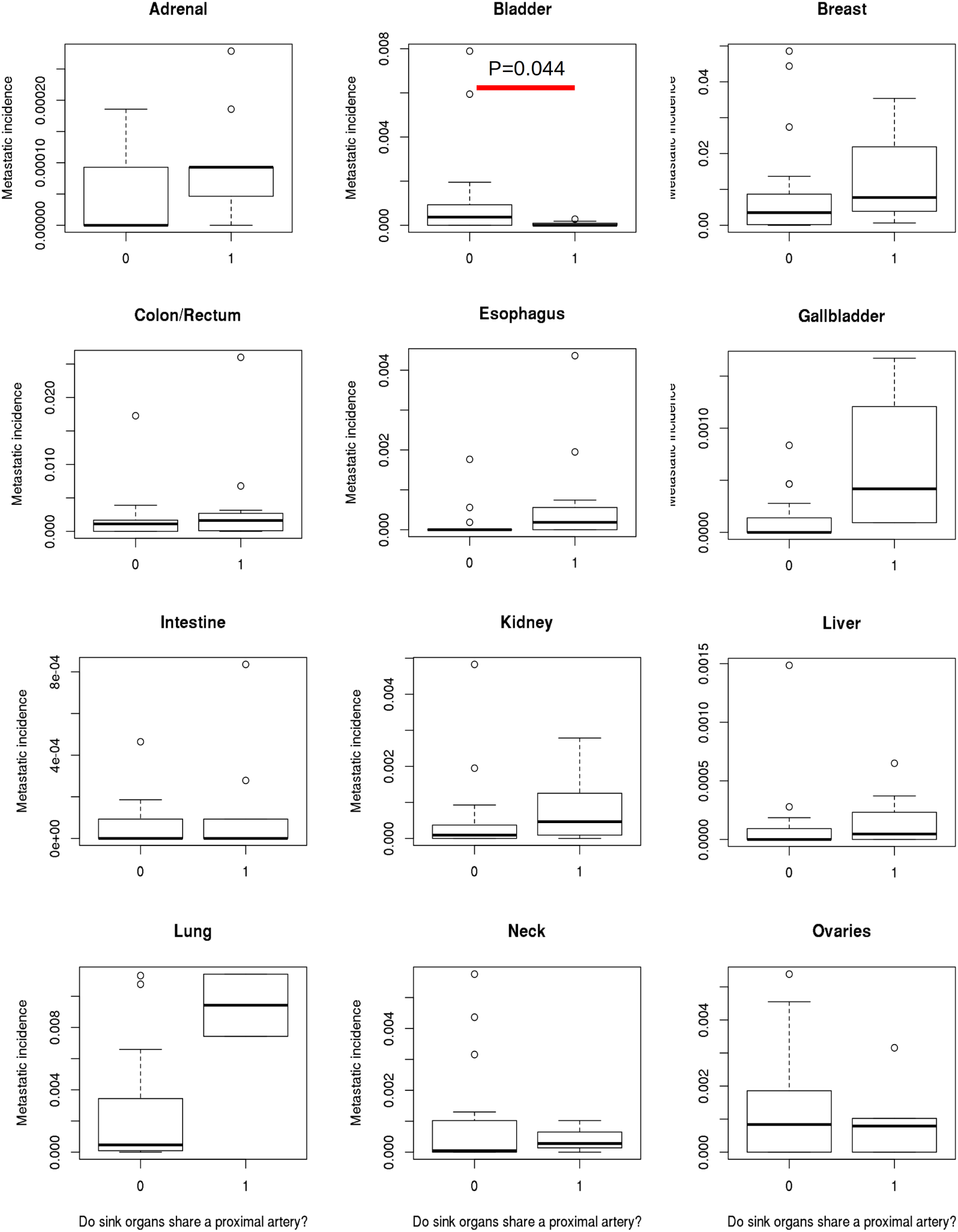
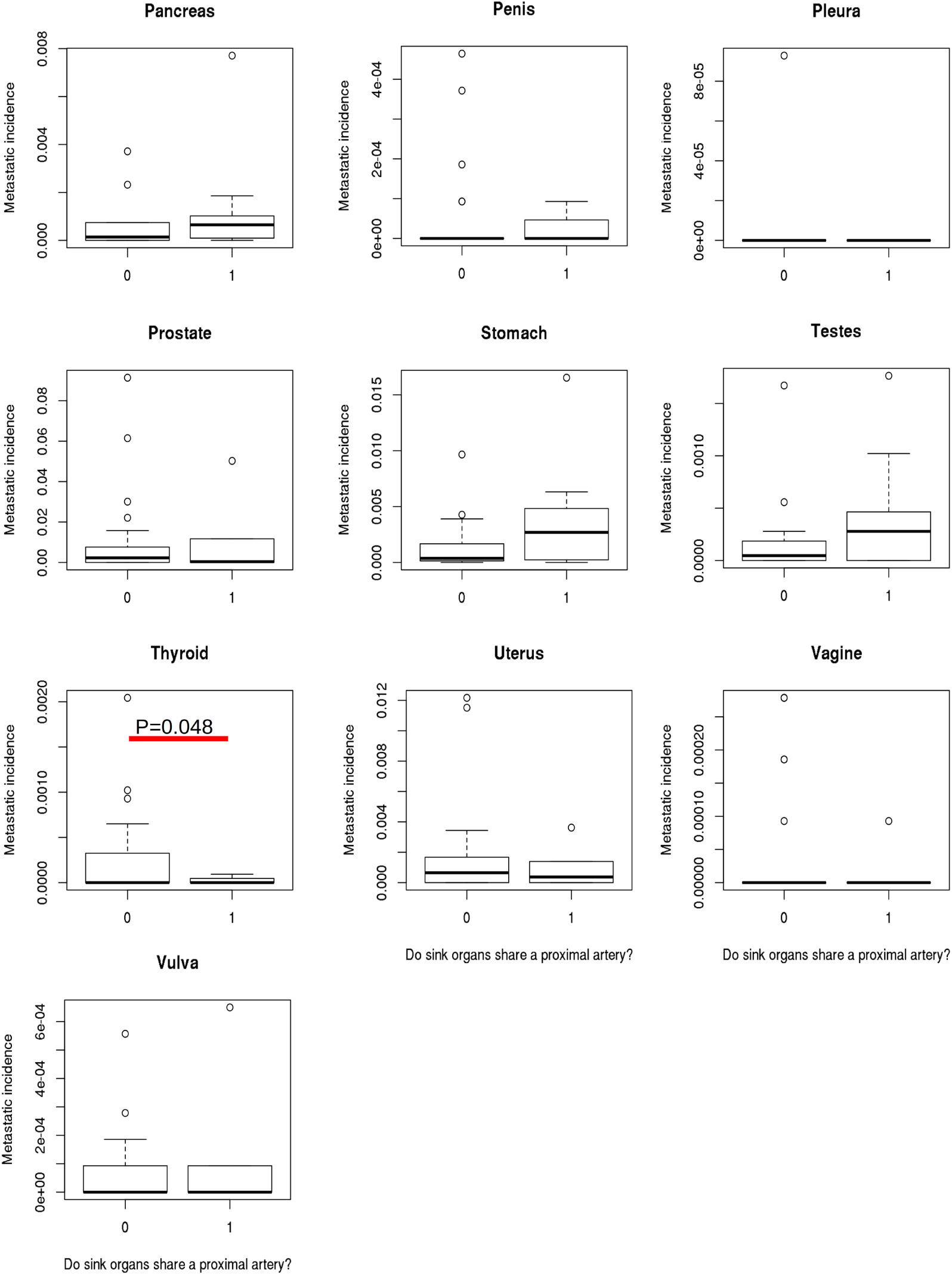
Each plot represents a primary tumor showing the relationship between broad sense incidence (BSI) and common blood supply (0=no, 1=yes) of acceptor organs with the corresponding source. Only statistical differences are shown (p<0.05). For each primary tumor, we tested (using the Student t-test) if organs sharing a vessel has an effect on the average metastatic incidence value. All statistical tests were evaluated under the same criteria (Type I error rate *α =* 0.05).

**Fig. S3.**
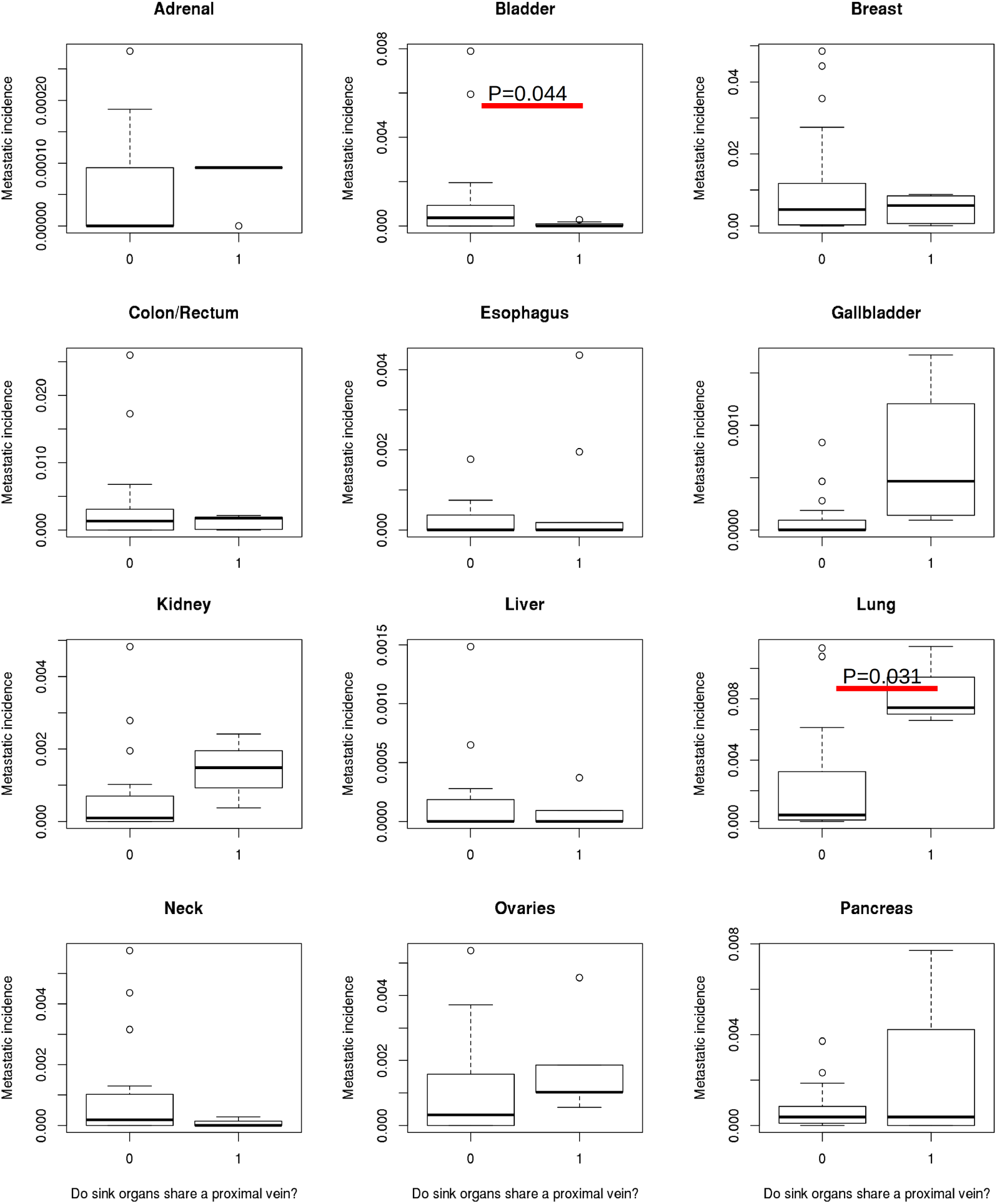
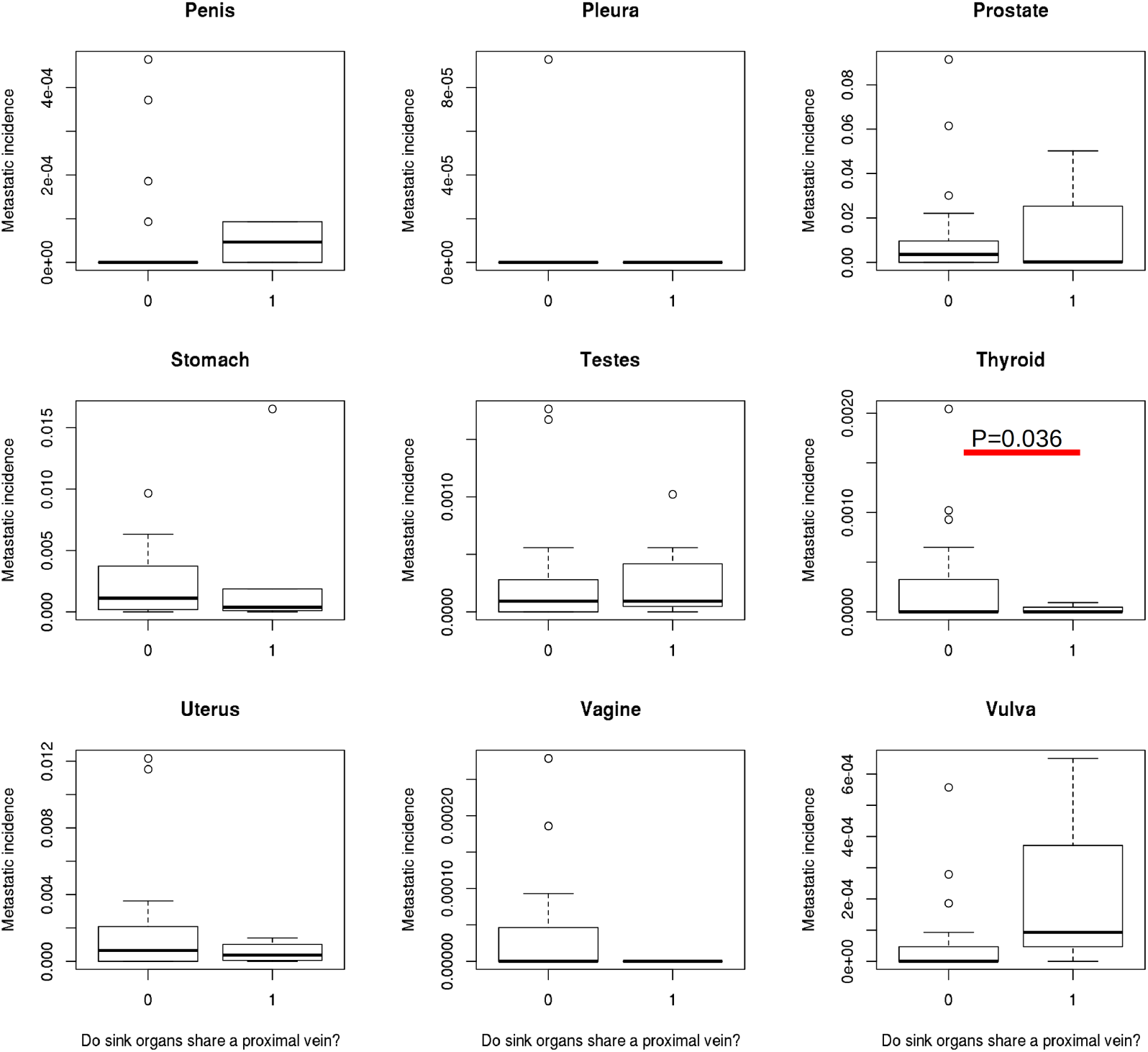
Each plot represents a primary tumor showing the relationship between broad sense incidence (BSI) and common blood drainage (0=no, 1=yes) of acceptor organs with the corresponding source. For each primary tumor, we tested (using the Student t-test) if organs sharing a vessel (arteries or veins) has an effect on the average metastatic incidence value. All statistical tests were evaluated under the same criteria (Type I error rate *α* = 0.05). Only statistical differences are shown (p<0.05).

**Table S1:** Summary of the different statistical fits applied to the probability distribution *P*[*k*] for the degrees *k* for source and sink organs. The truncated power law fits two coefficients: slope and cut-off; in this table only the slope estimation is shown.

| Source organs |  |  |  |  |  |
| --- | --- | --- | --- | --- | --- |
| Fit | Estimate | Std. error | $\Pr(> \vert t\vert )$ | $R^2$ | AIC |
| Exponential | 0.047 | 0.006 | $2.47 \times 10^{-6}$ | 0.898 | -17.585 |
| Power law | 0.257 | 0.072 | $2.9 \times 10^{-3}$ | 0.643 | 0.827 |
| Truncated power law | -0.357 | 0.055 | $1.48 \times 10^{-5}$ | 0.978 | -41.159 |
| Acceptor organs |  |  |  |  |  |
| Fit | Estimate | Std. error | $\Pr(> \vert t\vert )$ | $R^2$ | AIC |
| Exponential | 0.053 | 0.006 | $3.36 \times 10^{-8}$ | 0.923 | -26.932 |
| Power law | 0.308 | 0.061 | $7.71 \times 10^{-5}$ | 0.757 | -5.112 |
| Truncated power law | -0.263 | 0.081 | $4.6 \times 10^{-3}$ | 0.958 | -35.561 |

